# Learning of probabilistic punishment as a model of anxiety produces changes in action but not punishment encoding in the dmPFC and VTA

**DOI:** 10.1101/2022.03.29.486234

**Authors:** David S. Jacobs, Madeleine C. Allen, Junchol Park, Bita Moghaddam

## Abstract

Previously, we developed a novel model for anxiety during motivated behavior by training rats to perform a task where actions executed to obtain a reward were probabilistically punished and observed that after learning, neuronal activity in the ventral tegmental area (VTA) and dorsomedial prefrontal cortex (dmPFC) encode the relationship between action and punishment risk (Park & Moghaddam, 2017). Here we used male and female rats to expand on the previous work by focusing on neural changes in the dmPFC and VTA that were associated with the learning of probabilistic punishment, and with anxiolytic treatment with diazepam after learning. We find that adaptive neural responses of dmPFC and VTA during the learning of anxiogenic contingencies are independent from the punishment experience and occur primarily during the peri-action period. Our results further identify peri-action ramping of VTA neural activity, and VTA-dmPFC correlated activity, as potential markers for the anxiolytic properties of diazepam.

## Introduction

Anxiety is a debilitating symptom of most mental health disorders. Laboratory studies into the neural underpinnings of anxiety typically assess innate anxiety through exposure to an ambiguous context (Lezak et al., 2017). While this approach has provided valuable insight into the mechanisms which underlie acute fear and anxiety states, they do not address real-life situations where anxiety develops because actions are learned to be associated with a risk of an aversive outcome. An alternative approach to study this form of anxiety is to utilize uncertain punishment contingencies during action execution, which produce self-reported elevations in anxiety in humans and have been validated with anxiolytic treatment with benzodiazepines in rodents and non-human primates (Fischer et al., 2010; Schmitz & Grillon, 2012; Vogel et al., 1971). Recent procedures assessing reward and punishment learning simultaneously, have further indicated that learning of punishment contingencies may be critical to understanding behavioral and neuronal changes related to anxiety (Jean-Richard-Dit-Bressel, Lee, et al., 2021; Jean-Richard-dit-Bressel et al., 2019).

In our previous work (Park & Moghaddam, 2017), we assessed this mode of learned anxiety by developing a punishment risk task (PRT) in rats where actions executed to obtain reward conflicted with the presence of a low probability of harm (low intensity footshock). We observed that, after learning PRT, response time during risky actions became longer and more variable. Consistent with the relevance of PRT to anxiety, treatment with the anxiolytic drug diazepam diminished the impact of footshock probability on response time. To begin to understand the neural mechanisms that support PRT performance, we recorded from the dorsomedial prefrontal cortex (dmPFC) and the ventral tegmental area (VTA) after learning. The focus on these interconnected regions was because they have been implicated in anxiety (Balderston, Liu, et al., 2017; Balderston, Vytal, et al., 2017; Eysenck et al., 2007; Holmes & Wellman, 2009; Jacobs & Moghaddam, 2021; Park et al., 2016; Roberts, 2020) and are critical for execution of reward-guided actions (Balleine & Dickinson, 1998; Ellwood et al., 2017; Flores-Dourojeanni et al., 2021; Watabe-Uchida et al., 2017). We found that dmPFC and VTA neurons flexibly, and in coordination, encode the relationship between action and punishment. Here we sought to address two outstanding questions: 1) Do these brain regions “learn” punishment contingencies by changing their responses to the punishment or reward during learning? (2) After leaning, how are behavioral changes in response to anxiolytic treatment with diazepam encoded in these regions?

We used fiber photometry instead of single unit recording to assess the activity of neural population states in the dmPFC and VTA during PRT acquisition so that we could measure changes in the phasic neural response to the punishment (footshock) itself during task learning. After learning, we examined the effect of diazepam on dmPFC and VTA activity in correlation with behavior. We find that learning of probabilistic punishment contingencies is encoded by the VTA and dmPFC during action execution whereas punishment encoding remains unchanged. Moreover, anxiolytic treatment enhanced VTA action encoding and promoted correlated activity between the two regions without influencing dmPFC activity or encoding of the punishment.

## Methods

### Subjects

Thirteen adult Long-Evans (n=8) and Sprague-Dawley (n=5) rats, pair-housed on a reverse 12 h:12 h light/dark cycle, were used. All experimental procedures and behavioral testing were performed during the dark (active) cycle. Studies included both male (n=7) and female (n=6) rats. Animals were bred in house (n=8) or obtained from Charles River (n=5). All experimental procedures were approved by the OHSU Institutional Animal Use and Care Committee and were conducted in accordance with National Institutes of Health Guide for the Care and Use of Laboratory Animals.

### Surgery

#### Viral Infusion Surgery

Prior to task training, animals were injected with AAV8-hSyn-GCaMP6s-P2A-tdTomato (OHSU Vector Core, 5e13 ng/ml) to allow for pan-neuronal expression of fluorescent calcium indicator GCaMP6s and red fluorophore tdTomato. The coexpression of tdTomato allows for a motion artifact control signal to be used to correct GCaMP signals in rodents (Babayan et al., 2018; Matias et al., 2017; Menegas et al., 2018; Soares et al., 2016). Rats were anesthetized with isoflurane and placed in a stereotaxic apparatus. A small incision on the scalp was performed. The skull was cleaned and two craniotomies over the dmPFC and ipsilateral VTA were performed. A microinfusion syringe (Hamilton) was filled with virus and lowered into the dmPFC (AP +3.0, ML + 0.6, DV -3.3 mm from dura) or VTA (AP -5.4, ML + 0.6, DV -7.5 from dura) and injected into the brain at a volume of 700 nl at 50 nl/min using a syringe pump (World Instruments). Following infusion, virus was given 12-min to diffuse before the needle was slowly removed. The incision was then closed and covered with triple antibiotic. Animals were given 5 mg/kg of carpofen after surgery and allowed at least 1 week to recover from surgery.

#### Fiber Implant Surgery

After allowing at least seven weeks for virus expression, subjects were implanted with an optical fiber aimed at the dmPFC (dmPFC; AP +3.0, ML + 0.6, DV -3.3 mm from dura) and VTA (AP -5.4, ML + 0.6, DV -7.5 mm from dura) using surgical procedures outlined in the virus infusion section, with the exception that three additional bore holes were made for skull screws and fibers were secured to the skull using a light curing dental cement (Ivoclear Vivadent). Animals were given 1 week to recover from surgery before behavioral testing.

#### Initial Training & Punishment Risk Task (PRT)

The PRT follows previously published methods (Chowdhury et al., 2019; Park & Moghaddam, 2017). Rats were trained to make an instrumental response to receive a 45-mg sugar pellet (BioServe) under fixed ratio one schedule of reinforcement (FR1). The availability of the nosepoke for reinforcement was signaled by a 5-s tone. After at least three FR1 training sessions, PRT sessions began. PRT sessions consisted of three blocks of 30 trials each. The action-reward contingency remained constant, with one nose-poke resulting in one sugar pellet. However, there was a probability of receiving a punishment (300 ms electrical footshock of 0.3 mA) after the FR1 action, which increased over the blocks (0%, 6%, or 10% in blocks 1, 2 and 3, respectively). To minimize generalization of the action-punishment contingency, blocks were organized in an ascending footshock probability with 2-min timeouts between blocks. Punishment trials were pseudo-randomly assigned, with the first footshock occurring within the first five trials. All sessions were terminated if not completed in 180 mins.

#### Diazepam Treatment

Injectable diazepam (Pfizer/Hospira, Lake Forest, Il.) at a dose of 2.0 mg/kg or sterile saline (0.9% NaCl) were injected intraperitoneally 5 min before the start of the task. All injections given at a volume of *≤* 1.0 mL/kg.

#### Fiber Photometry System and Recording Procedures

Recordings were performed with a commercially available fiber photometry system, Neurophotometrics Model: FP3001 (NPM). Recording was accomplished by providing both 470 nm and 560 nm excitation light through the 400 um core patchcord to the dmPFC or VTA for GCaMP6s and tdTomato signals, respectively. Data were recorded using bonsai open source software (Lopes et al., 2015) and timestamps of behavioral events were collected by 5V TTL pulses that were relayed to an Arduino interfaced with bonsai software.

### Fiber Photometry Analysis

#### Peri-event analysis

Signals from the 465 (GCaMP6s) and 560 (tdTomato) streams were processed in Python (Version 3.7.4) using custom written scripts similar to previously published methods (Jacobs & Moghaddam, 2020). Briefly, 465 and 560 streams were low pass filtered at 3 Hz using a butterworth filter and subsequently broken up based on the start and end of a given trial. The 560 signal was fitted to the 465 using a least-squares first order polynomial and subtracted from 465 signal to yield the change in fluorescent activity (F/F= 465 signal - fitted 560 signal/ fitted 560 signal). Peri-event z-scores were computed by comparing the F/F after the behavioral action to the 4-2 sec baseline F/F prior to a given epoch. To investigate potential different neural responses to punishment execution vs anticipation, punished (i.e. shock) trials and unpunished trials were separated. Trials with a z-score value *>* 40 were excluded. From approximately 3,000 trials analyzed, this occurred on *<* 1% of trials.

#### Time Lagged Cross-Correlation Analysis

Cross correlation analysis has been used to identify networks from simultaneously measured fiber photometry signals (Sych et al., 2019). For rats with properly placed fibers in the dmPFC and VTA, correlations between photometry signals arising in the VTA and dmPFC were calculated for the peri-action, punishment, and peri-reward periods using the z-score normalized data. The following equation was used to normalize covariance scores for each time lag to achieve a correlation coefficient between -1 and 1:

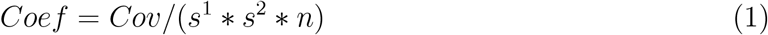

Where *Cov* is the covariance from the dot product of the signal for each timepoint, *s*^1^ and *s*^2^ are the standard deviation of the dmPFC and VTA streams, respectively, and *n* is the number of samples. An entire cross-correlations function was derived for each trial and epoch.

#### Comparison to Electrophysiology Results

Fiber photometry data for the third PRT session were compared to the average of the 50 msec binned single unit data (Park & Moghaddam, 2017, see Figure 4). This third PRT session corresponds to the session electrophysiology data were collected from. To overlay data from the two techniques, data were lowpass filtered at 3 Hz and photometry data were downsampled to 20 Hz (to match the 50 msec binning). Data from both streams were then min-max normalized between 0 and 1 at the corresponding cue and action+reward epochs.

To assess the similarity of the two signals, we performed a Pearson correlation analysis between the normalized single unit and fiber photometry data for cue or action+reward epochs at each risk block, as well as between randomly shuffled photometry signals with single unit response as a control. For significant Pearson correlations we performed cross correlation analysis (see above) to investigate if the photometry signal lagged behind electrophysiology given the slower kinetics of GCAMP6 compared to single unit approaches (Chen et al., 2013).

### Statistical Analysis

Trial completion was measured as the percentage of completed trials (of the 30 possible) for each block, while action latencies were defined as time from cue onset to action execution. Data were assessed through a repeated measures ANOVA or mixed effects model with factors risk block and session and *post-hoc* tests were performed using the bonferroni correction where appropriate. When only two groups were compared a paired t-test or Wilcoxon test was performed after checking normality assumption through the Shapiro-Wilk test. Statistical tests were performed using GraphPad Prism (Version 8) and utilized an *α* of .05.

To assess changes in neural activity, we utilized a permutation based approach as outlined in (Jean-Richard-dit-Bressel et al., 2020) using Python (Version 3). An average response for each subject for a given time point in the cue, action, or reward delivery period was compared to either the first PRT or saline session. For each time point a null distribution was generated by shuffling the data, randomly selecting the data into two groups, and calculating the mean difference between groups. This was done 1,000 times for each timepoint and a two-sided p-value was obtained by determining the percentage of times a value in the null distribution of mean differences was greater than or equal to the observed difference in the unshuffled data. To control for multiple comparisons we utilized a consecutive threshold approach based on the 3 Hz lowpass filter window (Jean-Richard-dit-Bressel et al., 2020; Pascoli et al., 2018), where a p value *<* .05 was required for 14 consecutive samples to be considered significant.

To assess correlated activity changes as a function of risk or session, we took the peak and 95% confidence interval for the overall cross correlation function. These values were compared by a two-way ANOVA with factors risk and session and utilized a *post-hoc* bonferroni correction. Tests were done using GraphPad Prism (Version 8) and utilized an *α* of .05.

Results for all statistical tests and corresponding figures can be found in Table 1 or supplemental figures.

**Table 1.**
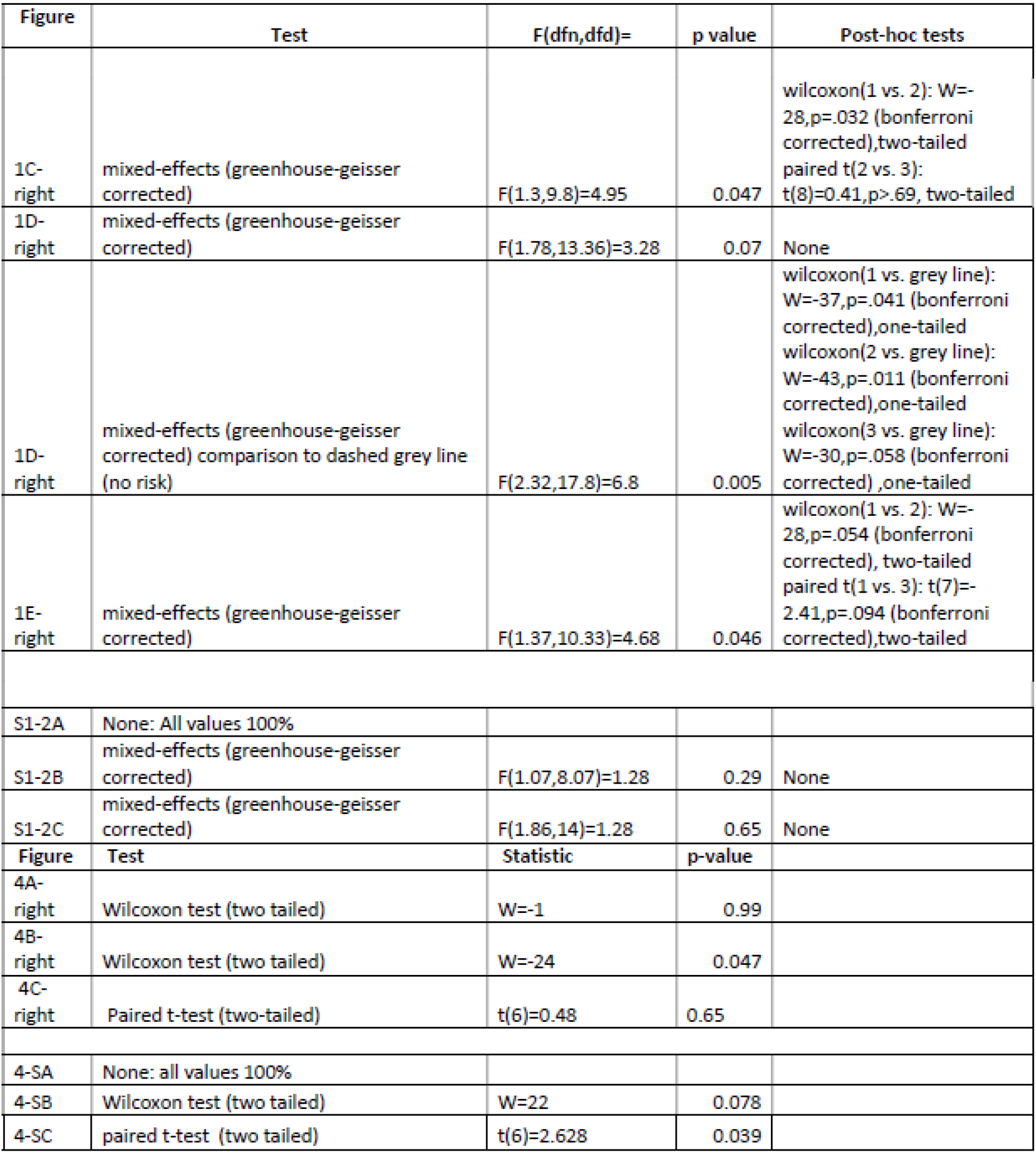
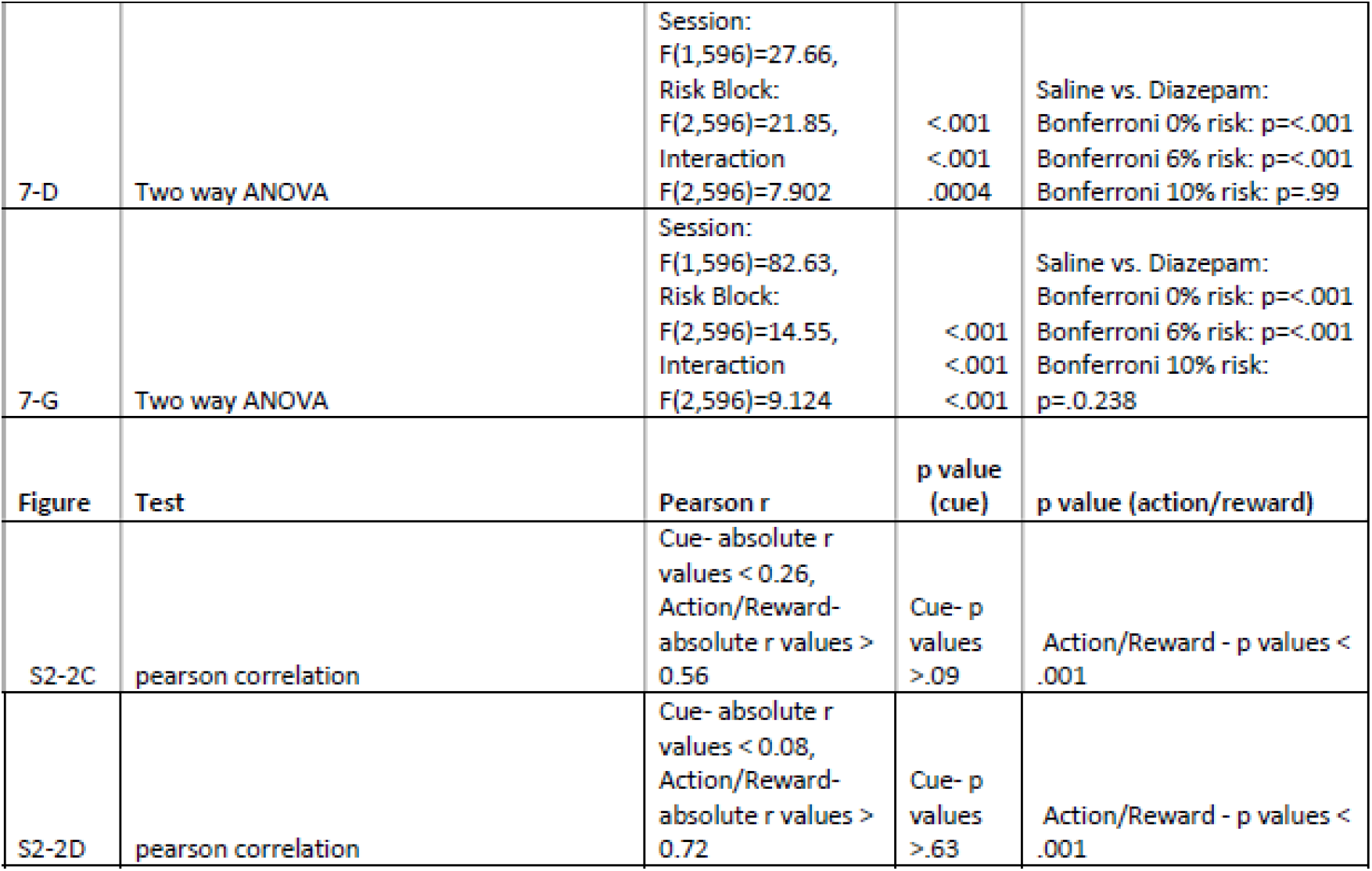
Statistical Results for Behavior and Correlation/Cross-Correlation Analyses Back to results.

### Excluded Data

One rat was excluded from behavioral and photometric analysis for Session 3 due to completion of only two trials. For the dmPFC, one session for a rat was excluded due to lack of timestamp collection and the first block of a session was excluded for a rat due to LED malfunction. Outliers from latency analysis were removed when a data point was *>* 5 SDs above the mean across all blocks. This removed one data point from analysis.

Four rats with VTA placement were excluded because fiber placements were too dorsal or ventral or GCaMP6s expression was not observed. Several rats did not complete all phases of the experiment due to lost fiber implants, leaving the final sample sizes as n=9 and n=7 for dmPFC in learning and diazepam treatment stages, respectively, and n=4 for VTA in learning and diazepam treatment stages.

### Histology & Imaging

Viral expression and fiber placements were verified after behavioral testing. Subjects were transcardially perfused with 0.01 M phosphate buffered saline (PBS) followed by 4% paraformaldehyde (PFA). Brains were removed and post-fixated in PFA for 24 h before being placed in 20% sucrose solution and stored at 4°C. Forty-µm brain slices were collected on a cryostat (Leica Microsystems) and preserved in 0.05% phosphate buffered azide. Brain slices were mounted to slides and cover slipped with Vectashield anti-fade mounting medium (Vector Labs). A Zeiss Apotome.2 microscope was used to image brain slices for GFP (Zeiss Filter set 38: 470-nm excitation/525-nm emission) and tdTomato (Zeiss Filter Set 43: 545-nm excitation/605-nm emission) to validate expression of both fluorophores in cells near the fiber tip. Fiber placement was determined by the brain slice demonstrating the most ventral fiber location.

Immunohistochemistry with a GFP antibody was used if a subject lacked virus expression to confirm the presence or absence of GCaMP6s. Brain slices were permeabilized in 3% BSA, 0.5% Triton X, and 5% Tween 80 dissolved in PBS + 0.05% sodium azide for 2 hr at room temperature. Slices were then incubated with rabbit antiserum against GFP (Abcam, Catalogue 6556, 1:500) diluted in PBS + Azide, 3% BSA + 0.1% Triton, and 1% Tween for 48 hr at 4°C. Slices were then washed in PBS + Azide, 3% BSA + 0.1% Triton + 1% Tween, three times for five minutes each. After this, slices were incubated with goat-antirabbit Alexa-488 (Abcam, Catlogue 1051G, 1:2000) diluted in PBS + Azide, 3% BSA + 0.1% Triton, and 1% Tween for 24 hr at 4°C and subsequently washed again as outlined above. Slices were then mounted to slides with Vectashield and imaged using the same procedures outlined above.

## Results

### Learning of probabilistic risk task (PRT)

To determine if learning of the PRT is associated with changes in behavior and neural activity, we recorded neural calcium activity in the dmPFC and VTA (Supp. Figure 1) using fiber photometry during the first three sessions of PRT training. Task training was as described before (Chowdhury et al., 2019; Park & Moghaddam, 2017). After rats learned to execute an action to receive a reward, a (varying) risk of shock was introduced contingent on the action (Figure 1A). The risk of footshock increased logarithmically from 0-10% during three 30-trial consecutive blocks (Figure 1B). During the first three sessions of PRT, learning was apparent by increases in the latency to execute to the risky action and decreases in trial completion (Figure 1C-D-left; Table 1). In Session 1, punishment risk was overgeneralized because there were modest increases in the latency to retrieve reward (Figure 1E). After Session 1, animals optimized their behavior, as suppression of action execution was observed specifically in risk blocks and did not change between Sessions 2 and 3 (Figure 1C-D-left; Table 1). Furthermore, reward retrieval increases from risk of footshock were attenuated in later sessions, though this effect only trended on significance after post-hoc bonferroni correction (Figure 1E-right). No behavioral changes were seen during PRT learning in the safe block, suggesting animals had learned to distinguish between blocks (Supp. Figure 2C-D-left).

**Figure 1:**
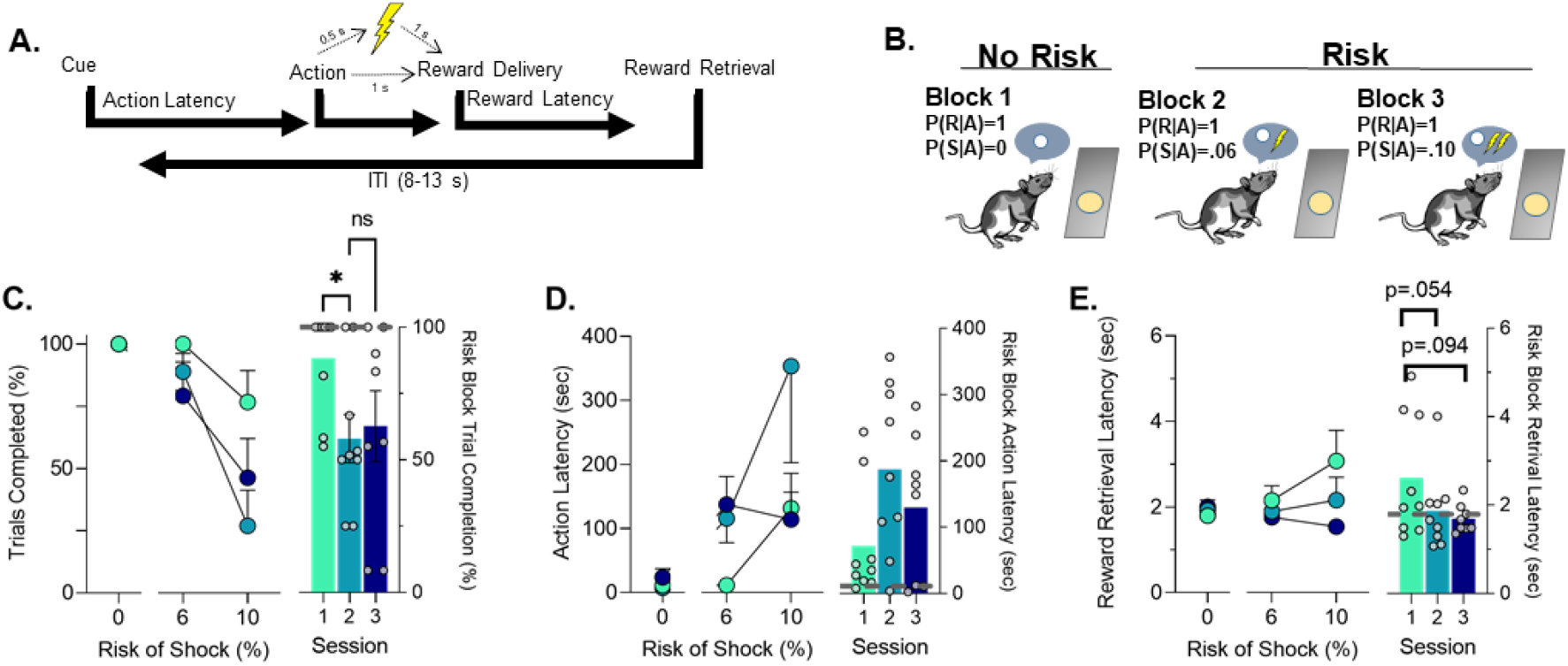
Schematic of PRT design and PRT behavior. **A**. Outline of trial structure in the PRT where action led to reward delivery with a varying risk of footshock. **B**. The multi-component schedule used to probabilistically punish the action with ascending risk of footshock. **C**. Trial completion over the first three sessions in each block and when comparing risk blocks over those sessions. **D**. Changes in latency to action completion over the first three sessions and specifically in risk blocks. **E**. Latency to retrieve the food reward over the first three sessions and specifically in risk blocks. Grey lines on bar graphs indicate the average for the safe block in Session 1 (i.e. before punishment was ever encountered). *p*<*.05, ns = not significant. *n* = 8-9 rats.

### dmPFC and VTA dynamically encode actions in PRT learning

Next we compared dmPFC and VTA neural activity in Sessions 1-3 to determine if neural responses to task events change in these regions during PRT learning. Data shown in Figure 2 and Figure 3 compare these responses in each block. For each session, we separated trials which resulted in footshock punishment from those that did not to determine if activity patterns were observed based on whether a trial was ultimately punished, or risky but ultimately safe as animals learned the task. Specifically, we were interested in whether a change in neural response to the actual punishment would occur during learning, or if changes in neural encoding of task events occurred when risk was present but footshock was not received. Figure 2 shows data from all three blocks excluding trials in block 2 and 3 where a footshock was received. In block 1 (the safe block) response to all task events remained the same during PRT learning. Session-by-session differences began to emerge in risk blocks. During 6% risk blocks, the initial modest peri-action inhibitory response decreased in both regions suggesting that action encoding is involved in learning the PRT (Figure 2B,E; Supp. Figure 3; Table 1). This peri-action response continued to be neutral or elevated during 10% risk blocks in the dmPFC and VTA, respectively. During the 10% risk blocks, learning of PRT was also associated with a transient increase in dmPFC peri-reward activation, and an increase in cue encoding in the VTA (Figure 2C and F, Supp. Figure 3). These results indicate that during learning of anxiogenic contingencies in the PRT both the VTA and dmPFC change their encoding of risky actions while the VTA also enhances its encoding of the cue.

**Figure 2:**
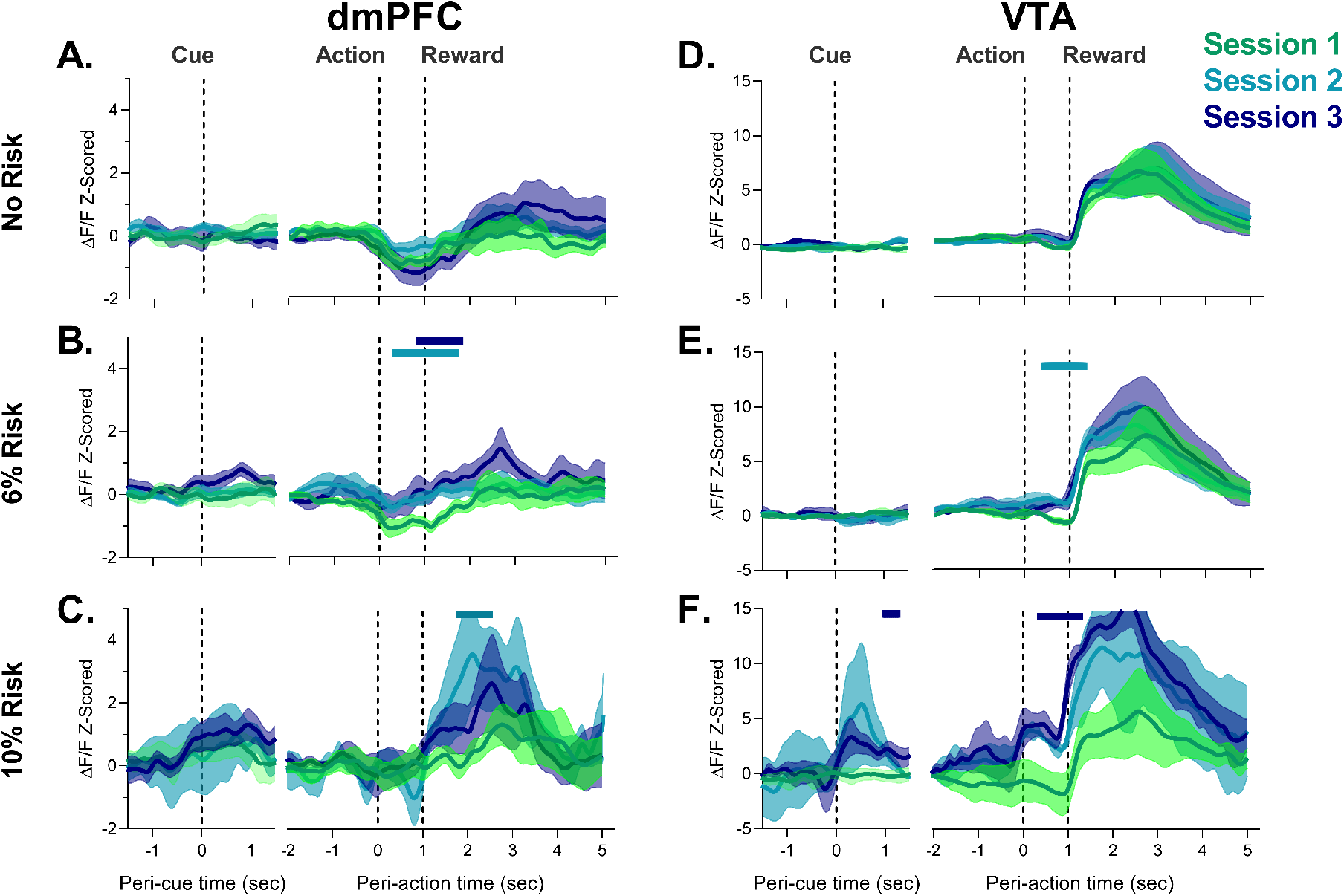
Neural responses to task events in unpunished trials by dmPFC and VTA during PRT learning. **A-C**. dmPFC responses to cue, action, and reward delivery for each block during the PRT task. Changes in dmPFC action encoding were observed when risk was present. **D-F**. VTA responses to cue, action, and reward delivery in the VTA. Changes in VTA action and cue and action responses were detected with task learning. Solid bars indicate significant differences from Session 1, where the color of the bar denotes the different Session. *n* = 3-9 rats, *n* = 2 rats for VTA Session 2 at 10% risk.

**Figure 3:**
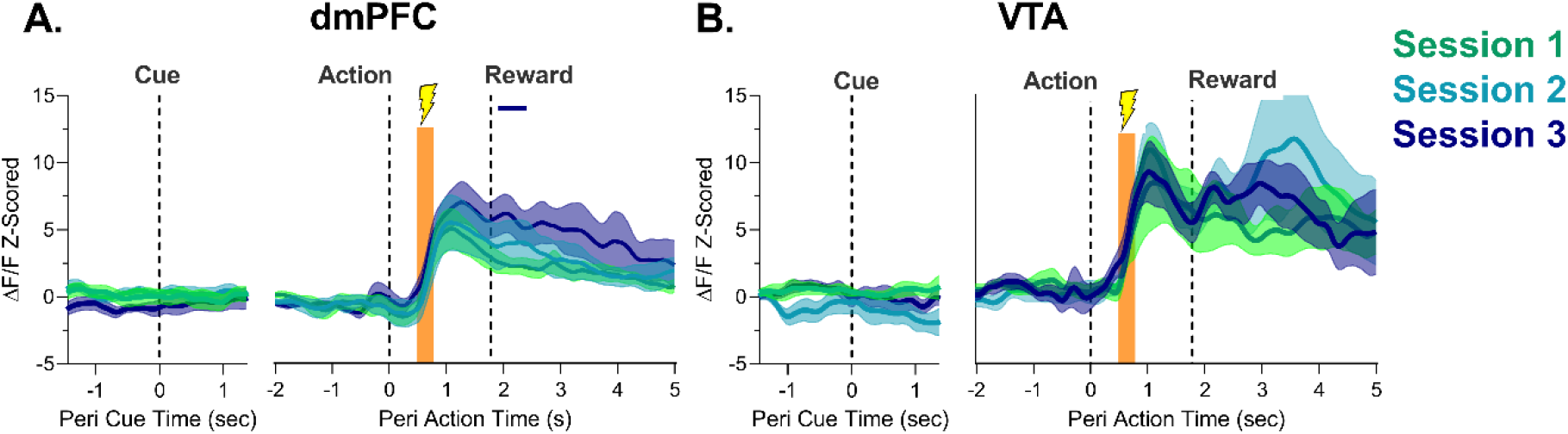
Neural responses to footshock punishment by dmPFC and VTA during PRT learning. **A**. The dmPFC demonstrated robust phasic increases in neural activity at the time of footshock administration over the three initial PRT sessions. **B**. Same as **A** but for the VTA. Orange bar indicates the period where footshock was administered. Solid lines indicate a significant difference from Session 1. *n* = 4-9 rats.

Figure 3 shows the session-by-session phasic response to footshock after action execution in blocks 2 and 3. Footshock produced a large increase in neuronal activity in dmPFC and VTA which reached its peak about 1 sec after action and 1 sec before reward delivery (Figure 3A-B). It is unlikely this increase is due to reward anticipation as in unpunished trials, the increase in activity did not peak until about 1.5-2 sec after action and comparing punished trials to unpunished trials revealed footshock related increases were seen significantly earlier than food delivery increases (Supp. Figure 5). The magnitude of this increase in the footshock/pre-reward period did not change in either region as animals learned the PRT (Supp. Figure 6), suggesting that learning of action-punishment contingency is not associated with increases or decreases in neuronal response to the punishment itself. Responses to task events at no risk or after footshock were not observed in animals where fiber placement was outside of the VTA with no GCaMP6s expression (Supp. Figure 7).

Decreases in the peri-action epoch in the dmPFC and the increase in response to reward delivery in the VTA are consistent with a large body of unit recording data (Mulder et al., 2003; Park & Moghaddam, 2017; Simon et al., 2015). To more directly establish a relationship between the present photometry responses and single unit activity at the corresponding session of PRT, we compared the current data with our single unit recording results (Park & Moghaddam, 2017). Both dmPFC and VTA signals were positively correlated at a higher level than when photometry data was randomly shuffled, mostly in the action and reward period (Supp. Figure 4). Relatedly, significant correlations were only observed in the action and reward period and not the cue period (Supp. Figure 4C-D,Table 1). The lack of detection of the faster (*<* 1 sec) cue responses by fiber photometry, particularly in some of the VTA data, compared to the more prolonged (*>* 1 sec) signal changes in action and reward periods may be related to GCaMP6s’ slower kinetics compare to unit approaches. Inspection of traces suggests phasic responses were occasionally slower to peak in photometry data (Supp. Figure 4 see E) and slower to begin to decay; the latter of which could also account for the low correlation seen in some of the faster epochs such as the cue. Overall these results suggest fiber photometry captures overall population encoding seen in these regions during action and reward in the PRT.

### Effect of diazepam on behavior and encoding of task events by dmPFC and VTA

After animals learned the PRT, we sought to determine if anxiolytic treatment changed the neural response to task events by the dmPFC and VTA during corresponding changes in behavior. Consistent with our previous results (Park & Moghaddam, 2017), a low dose of diazepam (2 mg/kg) attenuated the latency to execute an action when risk was present (Figure 4B) without influencing the number of trials completed (Figure 4A; Table 1). The effect of diazepam was most pronounced at the 6% risk block (block 2) potentially because the effectiveness of this low dose began to dissipate toward the end of block 3. Motoric effects of diazepam were noted in the safe block where a non-significant increase in action latency was observed in some subjects and an increase in latency to reward retrieval was observed (Supp. Figure 8 B,C; Table 1). These effects were, however, transient because action latency subsequently decreased and there was no increase in reward retrieval latency in risk blocks (Figure 4B-C-right; Table 1).

**Figure 4:**
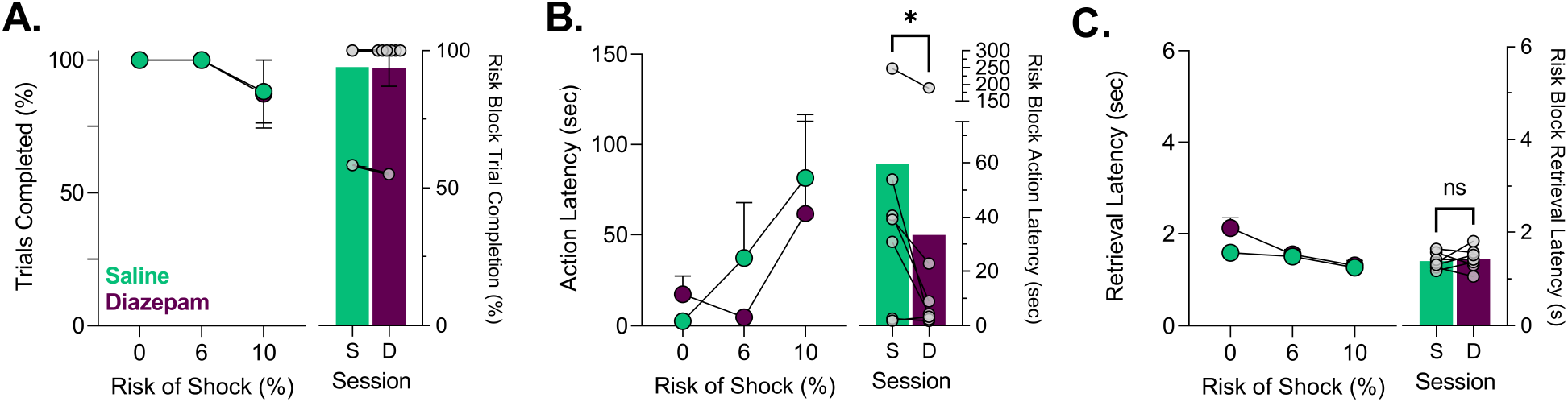
Effects of saline and diazepam (2 mg/kg) on PRT Behavior. **A**. Trial completion was unaffected by 2 mg/kg diazepam. **B**. Action latencies for trials where risk was present or absent. Diazepam significantly and consistently attenuated action latency increases seen from probabilistic punishment only when risk was present. **C**. Increases in reward retrieval latency which indicate motoric disruption from diazepam in block one dissipated in blocks 2 and 3 where risk was present. *p *<*.05, ns = not significant. *n* = 7 rats.

Diazepam influenced neural population activity differently in dmPFC and VTA. In the dmPFC, a reduction during the reward epoch was only observed in the safe block, and activity was not different from saline control levels in risk blocks across any epoch (Figure 5A-C). In the VTA, diazepam produced a ramping increase in population activity during the periaction epochs, just before action execution (Figure 5D-E, Supp. Figure 9) in no risk and 6% risk block (where diazepam had normalized behavior, Figure 4B). Similar VTA ramps using fiber photometry have been correlated with interoceptive goals (Guru et al., 2020). The sustained phasic increase in the VTA by diazepam may, therefore, provide a mechanism to explain its ability to enhance the likelihood of action execution under risk.

**Figure 5:**
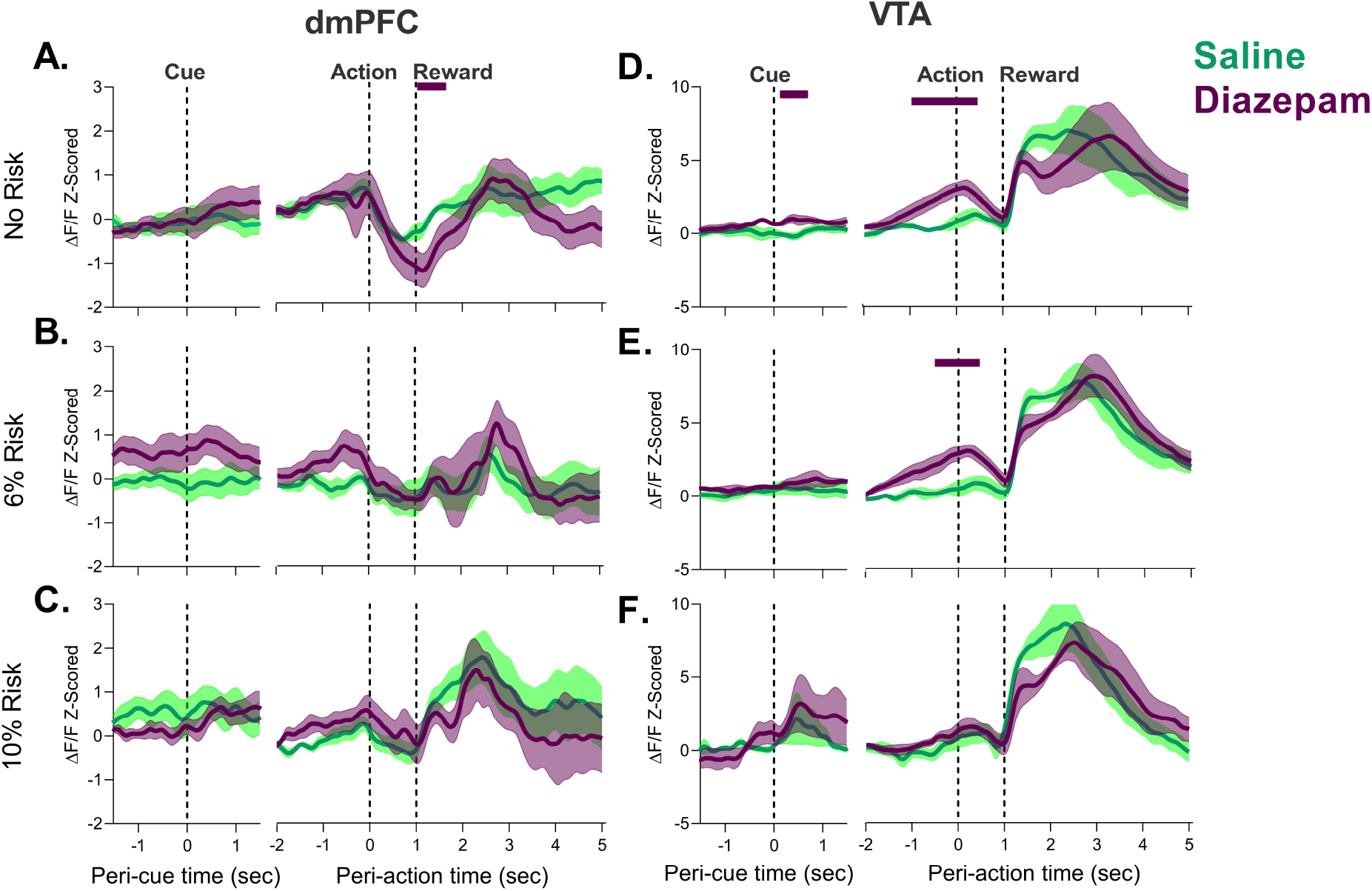
Effects of diazepam on neural activity during unpunished trials in the PRT in the dmPFC (left) and VTA (right). **A-C**. No effect of diazepam was observed during the cue or action epoch in the dmPFC, and a small but significant downward shift was seen following treatment early in the reward epoch in the safe block. **D-F**. Diazepam had no effect on neural activity during the reward period in the VTA. The peri-action activity was enhanced by diazepam until after action execution **D-E** but dissipated at high risk **F**. Solid lines above traces indicate significant differences from Saline at those timepoints. *n* = 4-7 rats.

Fiber photometry afforded the possibility to assess if diazepam’s anxiolytic effects may be related to changes in encoding of the punishment. Thus we separately analyzed the trials which resulted in footshock. Diazepam did not affect the neural response to footshock (Figure 6A-B). Both VTA and the dmPFC increased neural activity after footshock administration at comparable levels to that of saline (Supp. Figure 10). These results suggest that despite being an anxiolytic, diazepam does not change the encoding of the punishing stimulus by the dmPFC or VTA.

**Figure 6:**
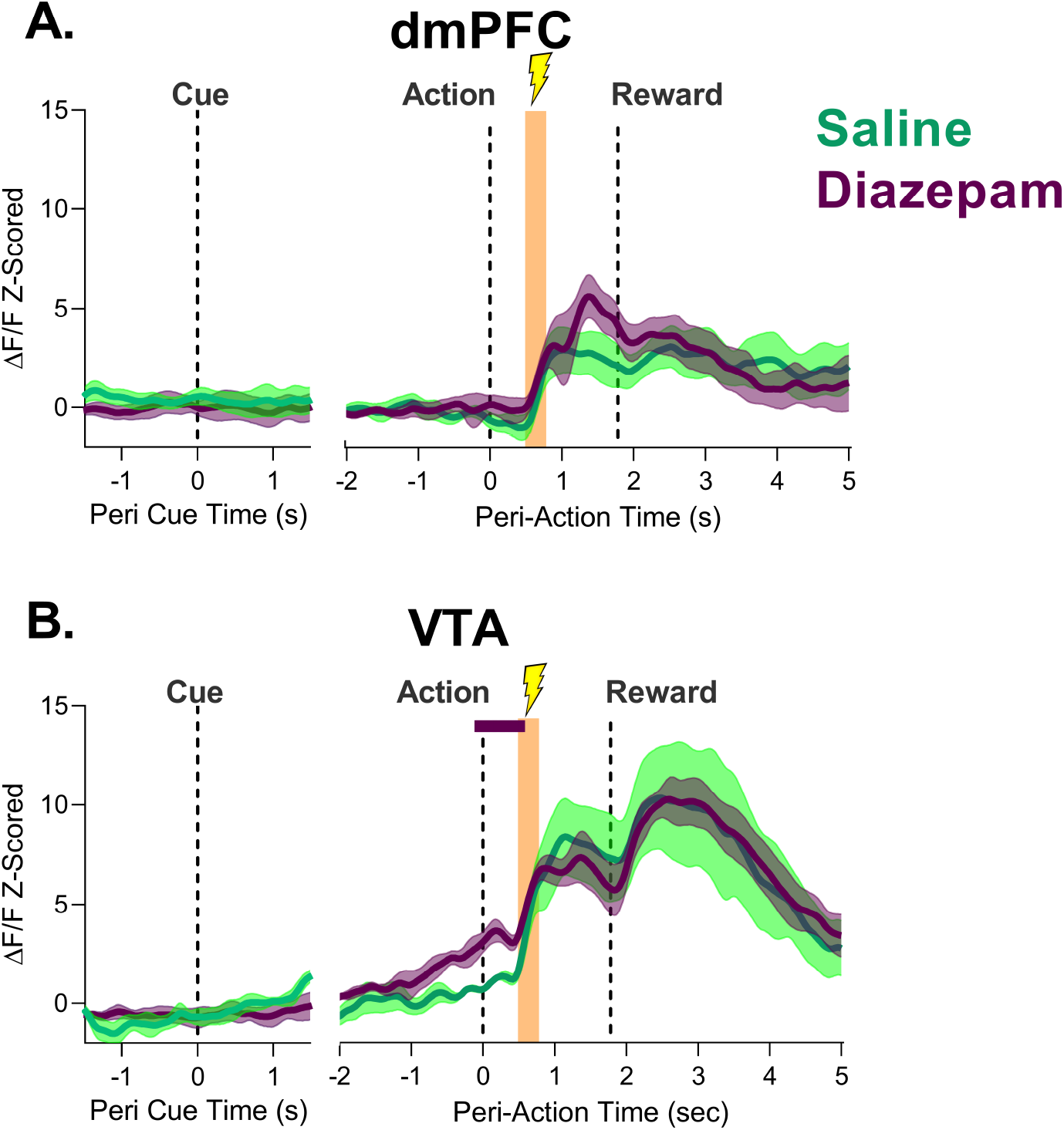
Effect of diazepam on neural response to footshock punishment in the dmPFC and VTA. **A**. dmPFC footshock responses were not different following diazepam treatment.**B**. VTA response to the footshock did not change with diazepam treatment. The only significant differences were observed during action execution, before the shock was administered. Orange bar indicates period of foot-shock administration. Solid lines indicate a significant difference from Saline. *n* = 4-7 rats.

Finally, because PFC co-activity with subcortical regions is critical for reward motivated behavior and implicated as a potential mechanism of anxiety (Balderston, Liu, et al., 2017; Balderston, Vytal, et al., 2017; Cornwell et al., 2017; Fujisawa & Buzsáki, 2011; Park & Moghaddam, 2017; Sartori & Singewald, 2019; J. Xu et al., 2019), we asked if diazepam influences the correlated activity of the dmPFC and VTA on a trial by trial basis. To assess this, we performed a cross correlation analysis for all trials after saline or diazepam treatment for the action and reward epochs, an approach which has been used with fiber photometry to identify networks as well as time differences in correlated signals (Sych et al., 2019). In saline sessions, correlated activity between the two regions was seen only in the reward epoch with risk. Diazepam increased the correlated activity of dmPFC and VTA during action epochs in the safe and 6% risk block. This increase in correlated activity, however, was not observed when the risk of footshock increased to 10% (Figure 7A-D, Table 1). In the reward epoch an increase in correlated activity was observed during all blocks (Figure 7E-H, Table 1). Across all analyses, peak correlations generally appeared with no time lag, except for the action period at high risk, where a 0.5 second VTA lead was observed. Of note, diazepam did not influence the correlated activity during footshock (Supp. Figure 11; Table 1). Taken together these results suggest that while diazepam does not change action and reward encoding separately in VTA and dmPFC, it influences the correlated activity between these regions during these epochs.

**Figure 7:**
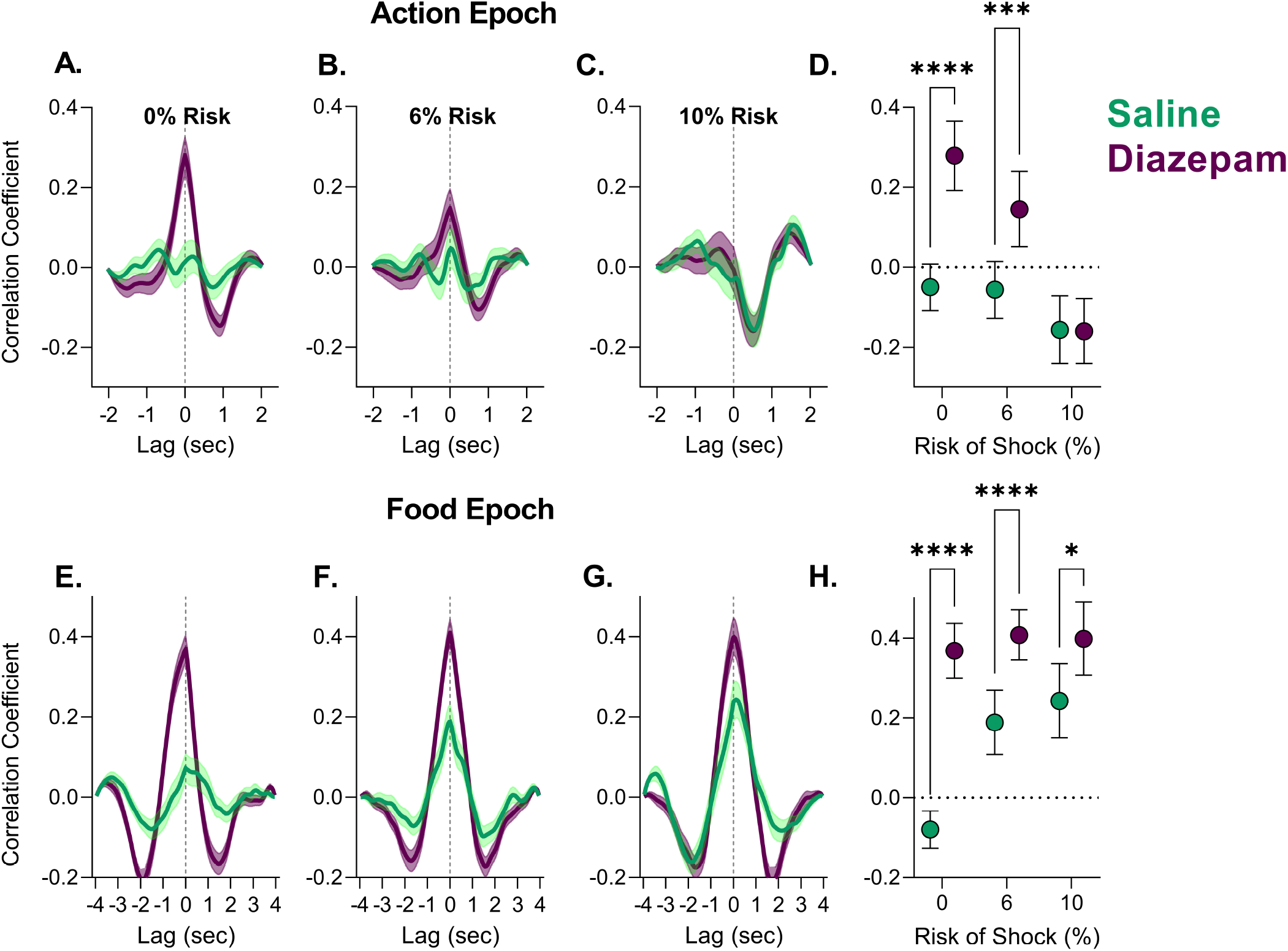
Correlated activity between the dmPFC and VTA during action and reward epochs in the PRT after saline or diazepam treatment. **A-C**. Correlated activity during action execution was enhanced by diazepam treatment in the safe block and the lower risk block. While correlated activity reached its lowest level at the highest risk block, regardless of treatment. **D**. Peak correlation coefficient values and 95% confidence interval for each cross-correlation function in **A-C. E-G**. Correlated activity was enhanced by diazepam during the reward epoch across all blocks. **H**. Peak correlation coefficient values and 95% confidence interval for each cross-correlation function in D-F. *p < .05, ****p < .001. *n* = 77-120 trials from 4 rats.

## Discussion

Learning and adapting to contingencies associated with rewarding or harmful outcomes are critical for survival. In the real world, these outcomes are not independent because actions executed to obtain a reward are often associated with a risk of an aversive outcome. This risk is learned over experience, and the perceived risk of punishment can engender anxiety-related states that will bias action selection. Using a novel task, our previous work demonstrated that the dmPFC and VTA dynamically encode risk of punishment during reward-motivated actions after rats have learned the punishment contingencies associated with these actions (Park & Moghaddam, 2017). The present study expands on these findings by focusing on the learning phase of the task, and the impact of diazepam after learning. We find that during punishment contingency learning, both VTA and dmPFC modify their response in the peri-action period but maintain similar responses to punishment and reward. Treatment with diazepam after learning did not influence the neural response to punishment in either region but it potentiated VTA action encoding and enhanced its correlated activity with the dmPFC. These data suggest that action, but not punishment or reward, encoding is critical to learning of punishment risk contingencies and the resulting state of anxiety.

### Punishment learning produces changes in action, but not punishment, encoding in the dmPFC and VTA

Learning of the punishment risk contingencies was apparent when the risk of footshock, which was initially overgeneralized to other aspects of the task, became selective for the risky action. The change in behavior was not related to dmPFC or VTA responsivity to the punishment itself as the magnitude of punishment encoding did not change in either region during learning. The large phasic response of these regions to footshock is consistent with previous literature implicating these regions in stress and pain responsiveness (Holly & Miczek, 2016; McKlveen et al., 2019) including our previous work showing that about 50-75% of spontaneously active PFC units respond to stress (Del Arco et al., 2020; Jackson & Moghaddam, 2006). Previous work has also shown that repeated exposure to the same stressor produces sensitization to some responses such as dopamine release in the prefrontal cortex (e.g. Gresch et al., 1994) and rapid desensitization of unit responses in the PFC (Jackson & Moghaddam, 2006). We, however, found that repeated exposure to footshock in the context of action contingent punishment did not produce a sensitized (or desensitized) response. This suggests that context matters in how PFC neurons respond to an aversive stimulus. While PFC neurons may adapt by reducing their response to a repeated punishment/stressor when stress presentation is certain or not contingent on an action, they appear to sustain the same level of activation when punishment presentation is uncertain and associated with an action.

Anxiogenic contingencies also did not influence the phasic response to reward by mPFC and VTA. In contrast to reward and punishment responses, peri-action responses changed in dmPFC and VTA as animals learned the PRT. This finding suggests that action encoding by the dmPFC and VTA can serve as a locus for punishment contingency learning during reward-guided behavior. This is an important observation because behavioral studies have shown that adaptation to reward and punishment is mostly associated with the learning of safe and/or dangerous contingencies rather than arousal from punishment or reward (Jean-Richard-Dit-Bressel, Lee, et al., 2021; Jean-Richard-dit-Bressel et al., 2019). The neural mechanisms which support reward-punishment contingency learning are thought to be diverse (McNaughton & Corr, 2004) with a recent study providing direct neural data for such processes (Jean-Richard-Dit-Bressel, Tran, et al., 2021). Our findings in the dmPFC and VTA implicate these regions in reward-punishment contingency learning and identify action encoding as a key behavioral event involved in this form of learning.

### Diazepam enhances VTA action encoding and VTA-dmPFC correlated activity without influencing punishment and reward encoding

Diazepam is a common anxiolytic drug that is used to validate anxiety assays and attenuates the action suppression seen from punishment risk in this task and others (Jacobs & Moghaddam, 2020; Liljequist & Engel, 1984; Park & Moghaddam, 2017). The mechanism for diazepam’s anxiolytic effects are poorly understood (Sartori & Singewald, 2019). One possibility is that diazepam itself attenuates responses to anxiogenic stimuli such as punishments. This mechanism was not supported by our data in that we did not see attenuation of punishment responsivity in the VTA or dmPFC after diazepam. An alternative explanation is that diazepam enhances responsivity to reward, which would consequently drive reinforced behavior under punishment risk. Again, our results did not support this explanation, as neural response to reward was similar across regions after drug treatment. Thus, diazepam has little impact on the processing of aversive or appetitive emotional stimuli in these regions. In contrast, diazepam influenced VTA activity during the peri-action epoch by producing a ramping of neural activity in the first two blocks before action execution. The so-called ‘ramping activity’ of VTA neurons has been previously associated with attentional tuning, movement kinetics, and distance to goals (Kremer et al., 2020; Totah et al., 2013). A recent study which elegantly characterized VTA ramps using fiber photometry found that these signals reflect interoceptive goals particularly when internal maps, and not external stimuli, are utilized to process reward proximity (Guru et al., 2020). One possible explanation for our results is that anxiogenic contingencies render subjects more attentive to stimuli and external conditions. Thus, diazepam’s production of VTA ramping activity may be a mechanism to direct attentional processes to serve internalized goal driven states, and ultimately, more efficient reward seeking behavior. This interpretation is also in line with studies which assess the cognitive effects of diazepam in humans, as diazepam has been shown to attenuate vigilant-avoidant patterns of emotional attention to fearful stimuli (Pringle et al., 2016).

While diazepam had negligible effects in the dmPFC, it influenced correlated activity between this region and VTA during action epochs in the safe and 6% risk block. This pattern of increased correlation of activity corresponded to the ramping signal in the VTA suggesting that a potential downstream effect of VTA’s response to diazepam is to increase its correlative activity with the dmPFC. Correlated activity between PFC units or local field potentials (LFPs) with subcortical regions including the VTA has been observed during other goal-directed behaviors including PRT (Fujisawa & Buzsáki, 2011; Mininni et al., 2018; Park & Moghaddam, 2017; J. Xu et al., 2019). Thus while diazepam may not change the activity of dmPFC neurons, it may produce some of its effects by altering correlative activity between dmPFC and VTA. Future investigations using unit and LFP recordings will be needed to better characterize the mechanism for this change in correlated activity under anxiety.

Effects of diazepam on behavior and VTA dissipated by the last and the highest risk block, when anxiety is presumably highest. Diazepam’s half-life is about 1 hr (Friedman et al., 1986) and it is possible that this observation is due to lower receptor occupancy by the third (highest risk) block. Another possibility is that diazepam may function in different capacities when threats are either distal or lower in likelihood, i.e. blocks 1 and 2, compared to when the threat probability is higher or certain. Future studies could address these possibilities through longer acting anxiolytics or temporally specific manipulations to systematically disrupt the transient and longer lasting effects from diazepam observed here.

### Relationship of fiber photometry data to previous electrophysiology findings

This work provides a substantial expansion of our prior paper by utilizing fiber photometry to understanding neural activity during learning because it allowed us to measure neural responses during footshock punishment. While fiber photometry signals are not a direct measure of spiking seen in single unit recording, and additional studies investigating its exact relation to single-unit approaches are nascent (see Legaria et al., 2021; Sych et al., 2019; H. Xu et al., 2021), our observations that dmPFC action-related activity was most sensitive to risk of punishment and the VTA developed phasic increases at the time of action execution is consistent with unit recordings measured in Park and Moghaddam, 2017. Similarly, we observed a large reward response in the VTA consistent with previous electrophysiology findings (Park & Moghaddam, 2017; Watabe-Uchida et al., 2017). Taken together these results provide support that calcium activity of dmPFC and VTA neurons through fiber photometry shows similarities to the overall population responses seen from in vivo unit recording in this task.

### Conclusion

Assessing how the brain encodes reward-directed actions when conflicted by punishment probability is a novel approach to model learned anxiety. Using an animal model to study this form of learning, we find that dmPFC and VTA selectively adapt their action encoding response during the learning of anxiogenic contingencies without modifying their response to the punishment. We also identify peri-action ramping of VTA activity as a potential marker for anxiolytic properties of diazepam in this model. The VTA ramping signal has been linked to preparatory attention and goal driven states. Potentiation of this response may lead to attentional disengagement from harm probability and provide a mechanism for how diazepam influences action execution under anxiety.

## Acknowledgements

The authors thank Michelle Kielhold for assistance with immunostaining and Alina Bogachuk for help with microscope imaging. D.S.J. is a recipient of ARCS foundation scholar award.

## Supplemental Figures

**Supp. Figure 1:**
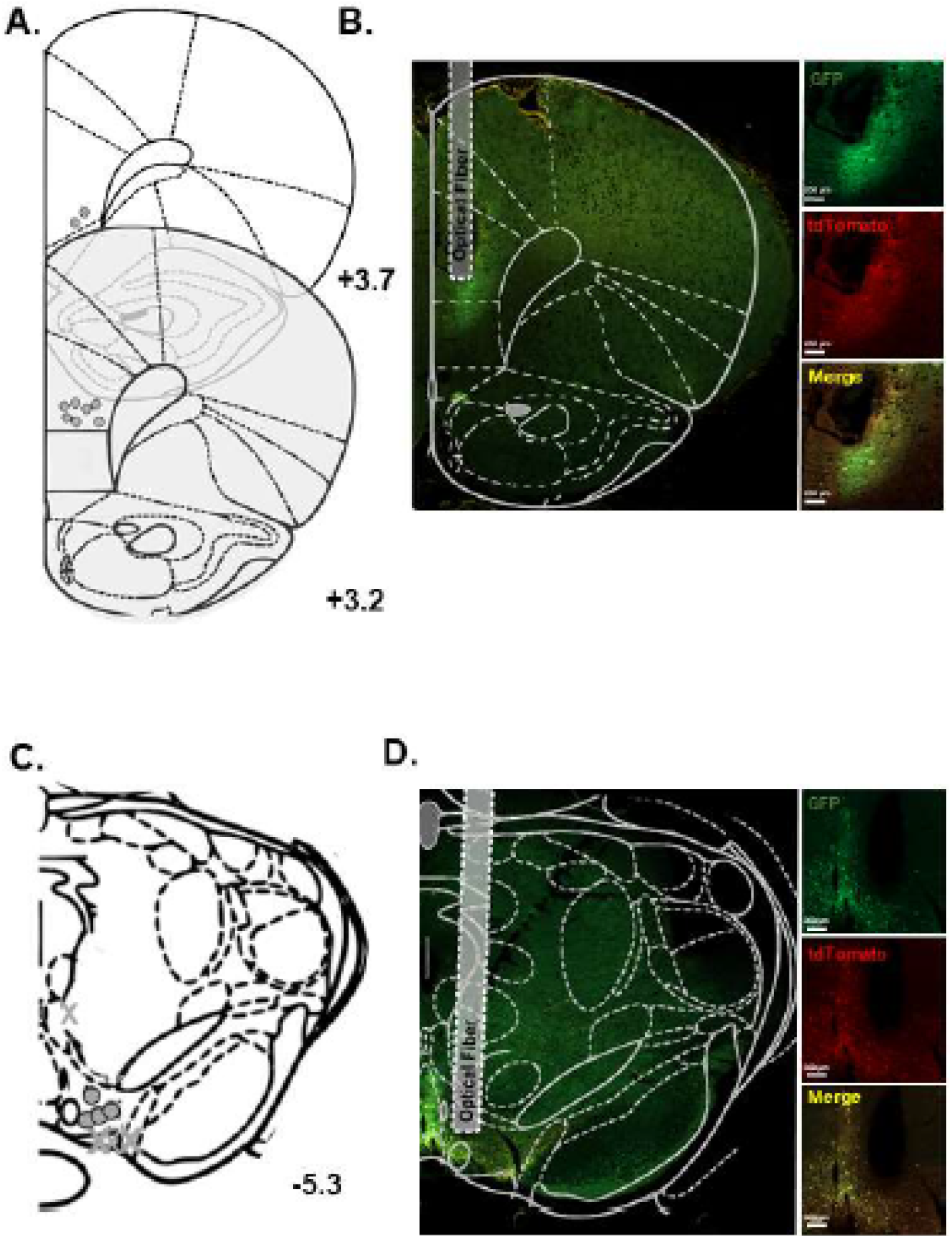
Hit map and representative image of GFP-GCaMP6s and tdTomato expression in the dmPFC and VTA. **A**. Fiber locations for individual subjects for the dmPFC. **B**. Representative image demonstrating expression of both GFP-GCaMP and tdTomato around the fiber tip in the dmPFC. **C**. Fiber locations for individual subjects for the VTA. X indicates placement of excluded subjects. **D**. Representative image demonstrating expression of both GFP-GCaMP and tdTomato around the fiber tip in the VTA. Scale bar= 200 µm. Brain outlines adapted from (Paxinos & Watson, 1998). Back to results.

**Supp. Figure 2:**
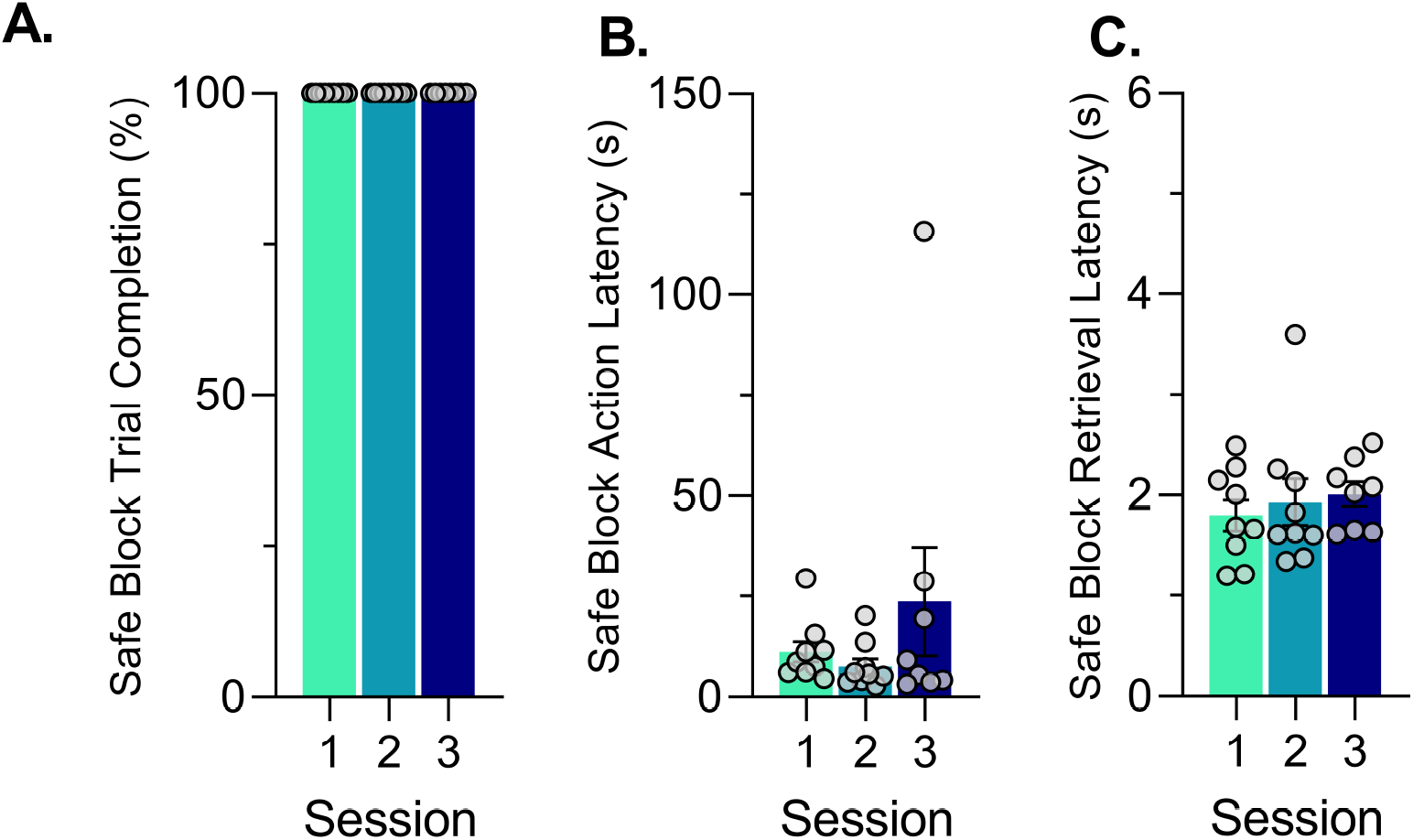
Average and individual (grey dots) behavior during the safe (0% risk) block for the first three learning sessions. **A**. Percentage of completed trials in the safe block. **B**. Latency to perform the action for the safe block. **C**. Latency to retrieve the food pellet in the safe block. Green - Session 1, Light Blue-Session 2, Purple – Session 3. Back to results.

**Supp. Figure 3:**
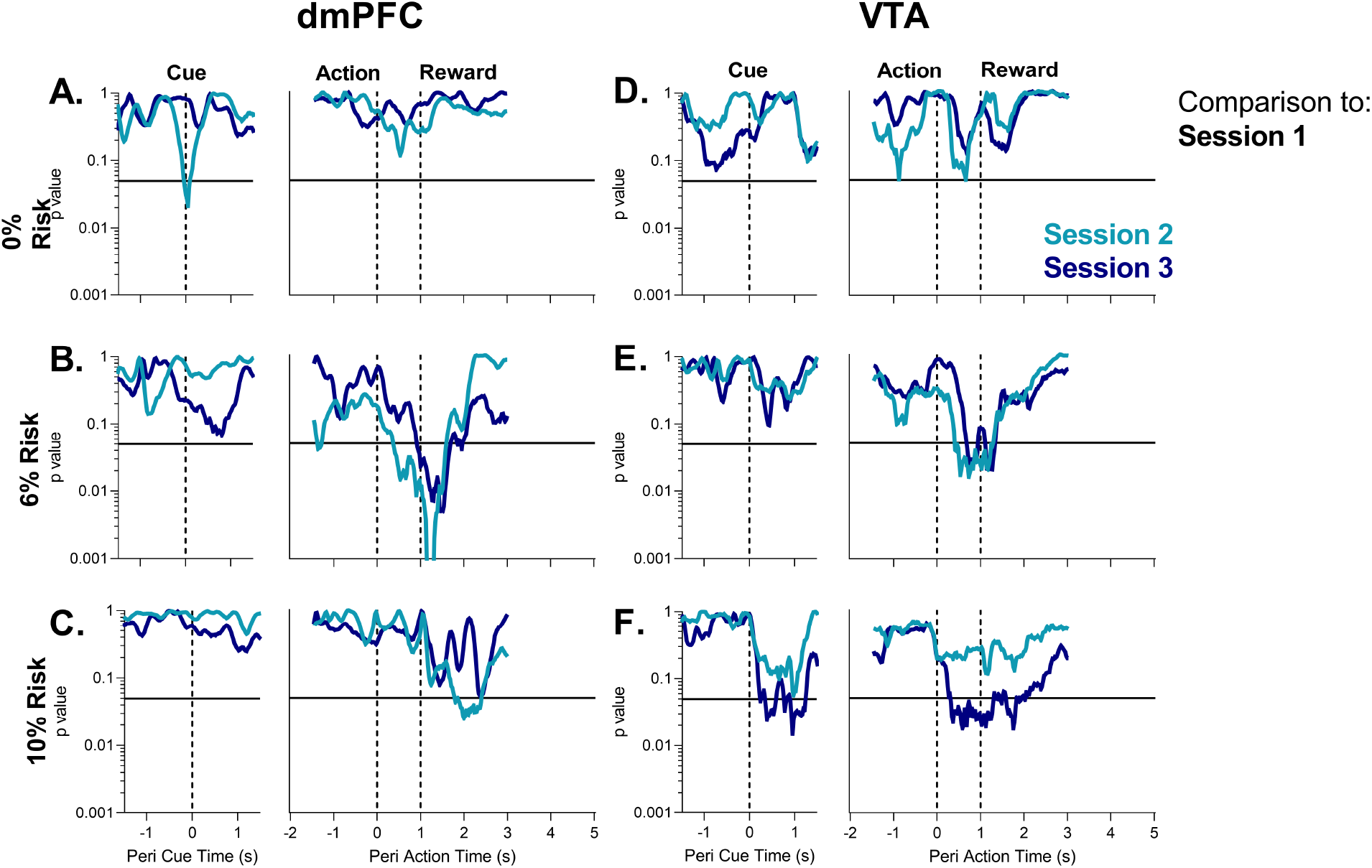
Permutation test results for recordings in the dmPFC and VTA for unpunished trials during PRT learning. **A-C**. dmPFC p-value for tests in the cue, action, and reward delivery periods for each block during the PRT task. **D-F**. VTA p-value for tests in the cue, action, and reward delivery periods for each block during the PRT task. All comparisons are to the first PRT session (Session 1). Light Blue-Session 2, Purple – Session 3. Solid black line indicates p =.05.Back to results.

**Supp. Figure 4:**
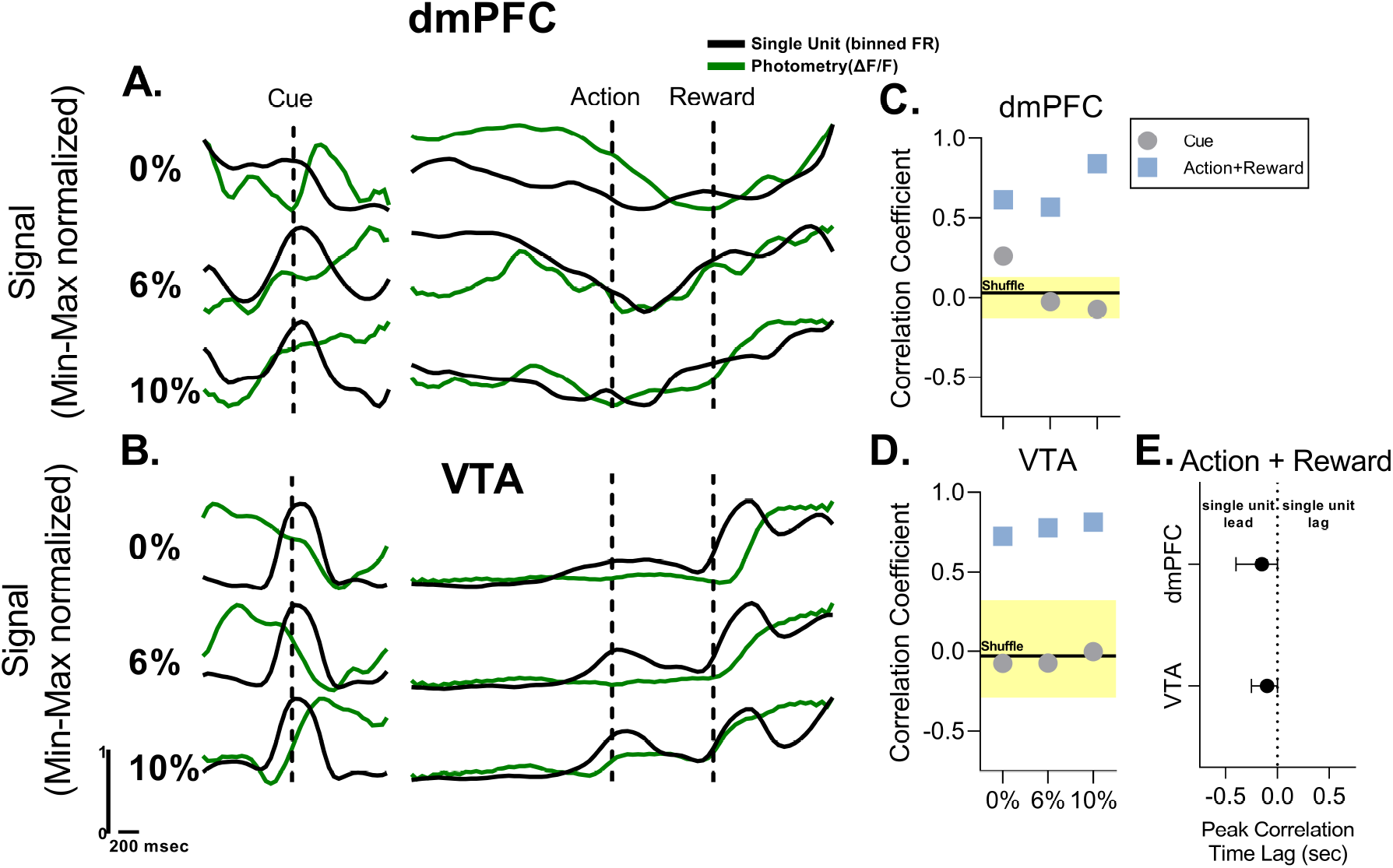
Comparison between average dmPFC and VTA fiber photometry (green trace -current study) and electrophysiological single unit recordings (black trace- (Park & Moghaddam, 2017)) during the corresponding session and epochs in the PRT. Data are min-max normalized to adjust for differences in magnitude of the signals. **A**. Changes in signal for photometry and single unit recordings in cue, action, and reward periods for the dmPFC. **B**. Changes in signal for photometry and single unit recordings in cue, action, and reward periods for the VTA. Single unit data reflect the average of putative DA and non-DA units. **C**. dmPFC correlation coefficient for cue and action+reward periods for comparison of single unit and fiber photometry. Black line and yellow shading reflects mean and 95% CI for correlation coefficient after random shuffling of the photometry signal. **D**. VTA correlation coefficient for cue and action+reward periods for comparison of single unit and fiber photometry. Black line and yellow shading reflects mean and 95% CI for correlation coefficient after random shuffling of the photometry signal. **E**. Lag for peak correlation coefficients discovered after time-lagged cross correlation in the dmPFC and VTA. Data reflect the mean of all three blocks with bars representing range of the data. Back to results.

**Supp. Figure 5:**
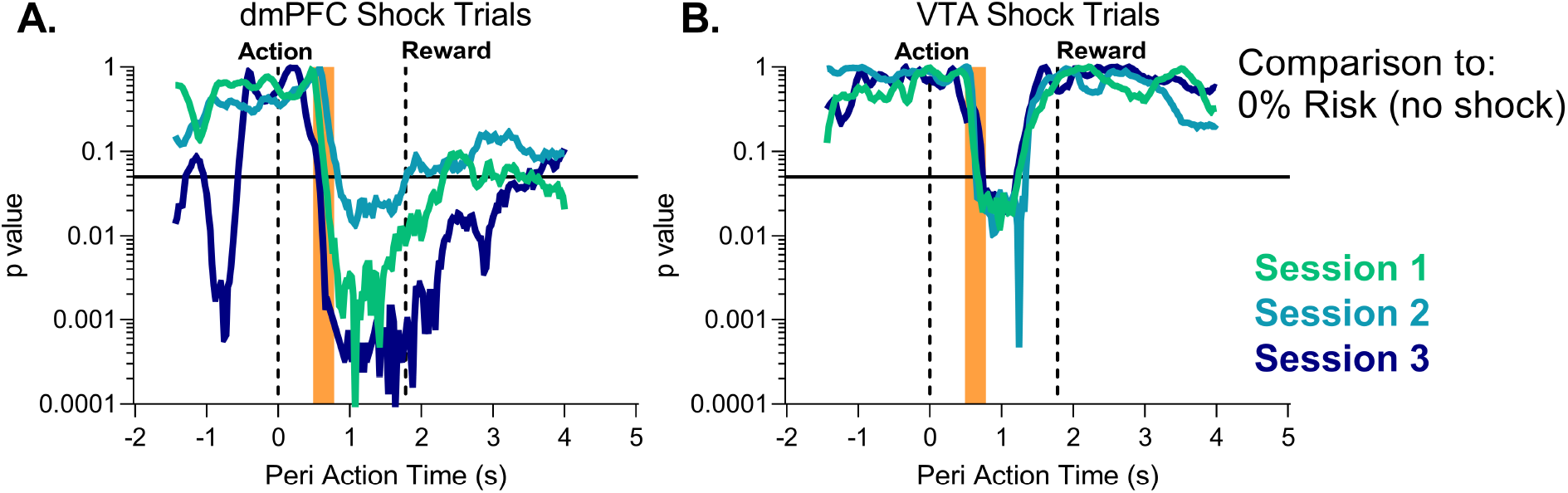
Permutation test results for recordings in the dmPFC and VTA for punished trials during PRT learning. **A**. dmPFC p-value for tests in the action, footshock (orange bar), and reward delivery periods for the first three session in the PRT task. **B**. VTA p-value for tests in the action, footshock (orange bar), and reward delivery periods for the first three session in the PRT task. All comparisons are to the corresponding time points in the 0% risk block (no footshock). Green, Session 1, Light Blue-Session 2, Purple – Session 3. Solid black line indicates p =.05.Back to results.

**Supp. Figure 6:**
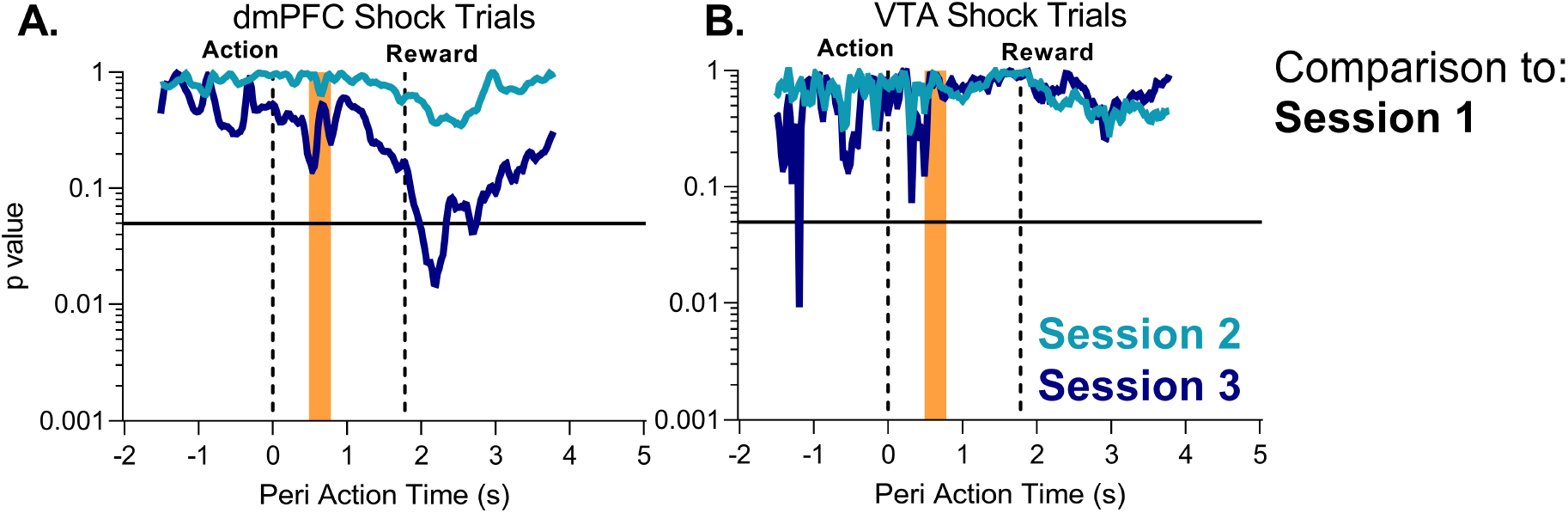
Permutation test results for recordings in the dmPFC and VTA for punished trials during PRT learning. **A**. ddmPFC p-value for tests in the action, footshock (orange bar), and reward delivery periods for the first three session in the PRT task. **B**. VTA p-value for tests in the action, footshock (orange bar), and reward delivery periods for the first three session in the PRT task. All comparisons are to the first PRT session. Light Blue-Session 2, Purple – Session 3. Solid black line indicates p =.05.Back to results.

**Supp. Figure 7:**
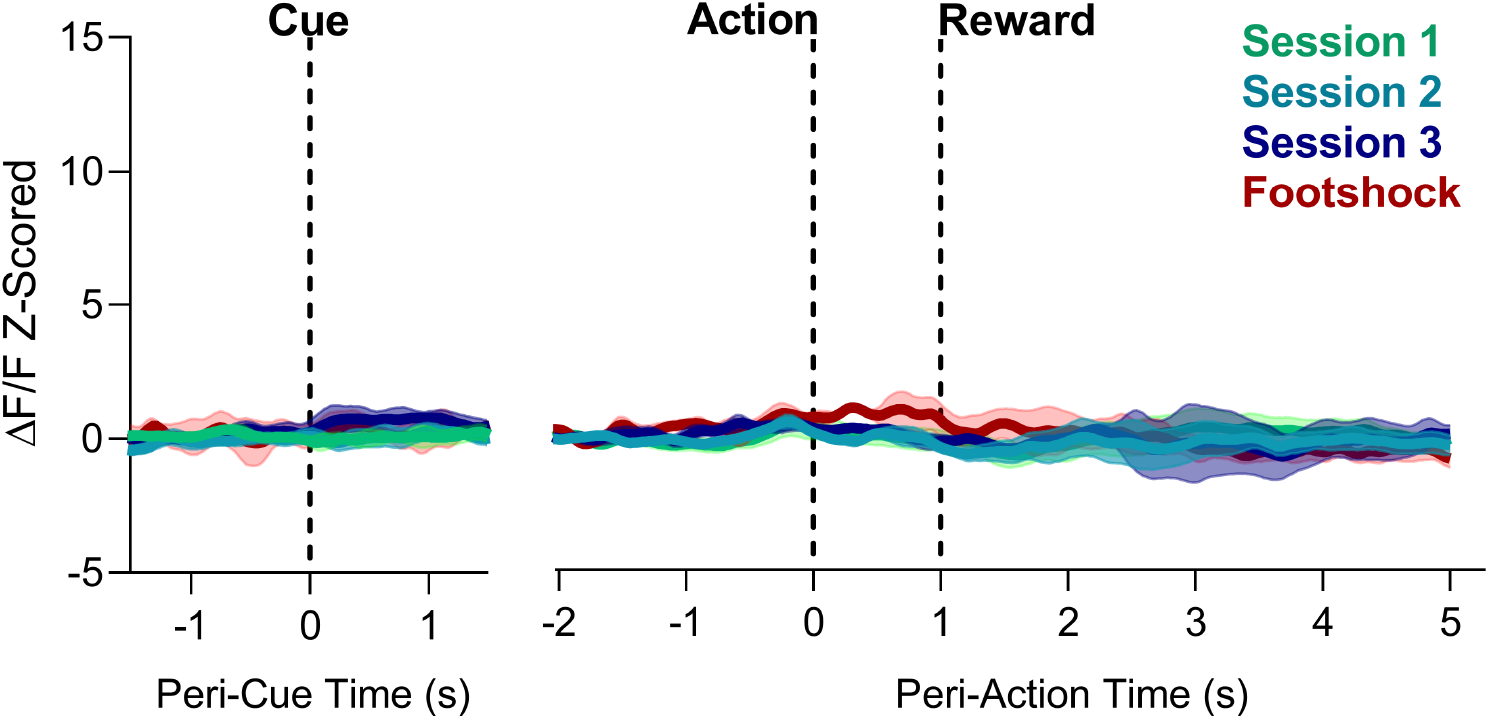
VTA fiber photometry traces for unpunished (0% risk block) and punished (Footshock) trials during PRT learning in rats with misplaced fibers or no GCaMP6s expression. Large responses to the footshock or reward delivery were not observed. *n* = 3-4 rats. Back to results.

**Supp. Figure 8:**
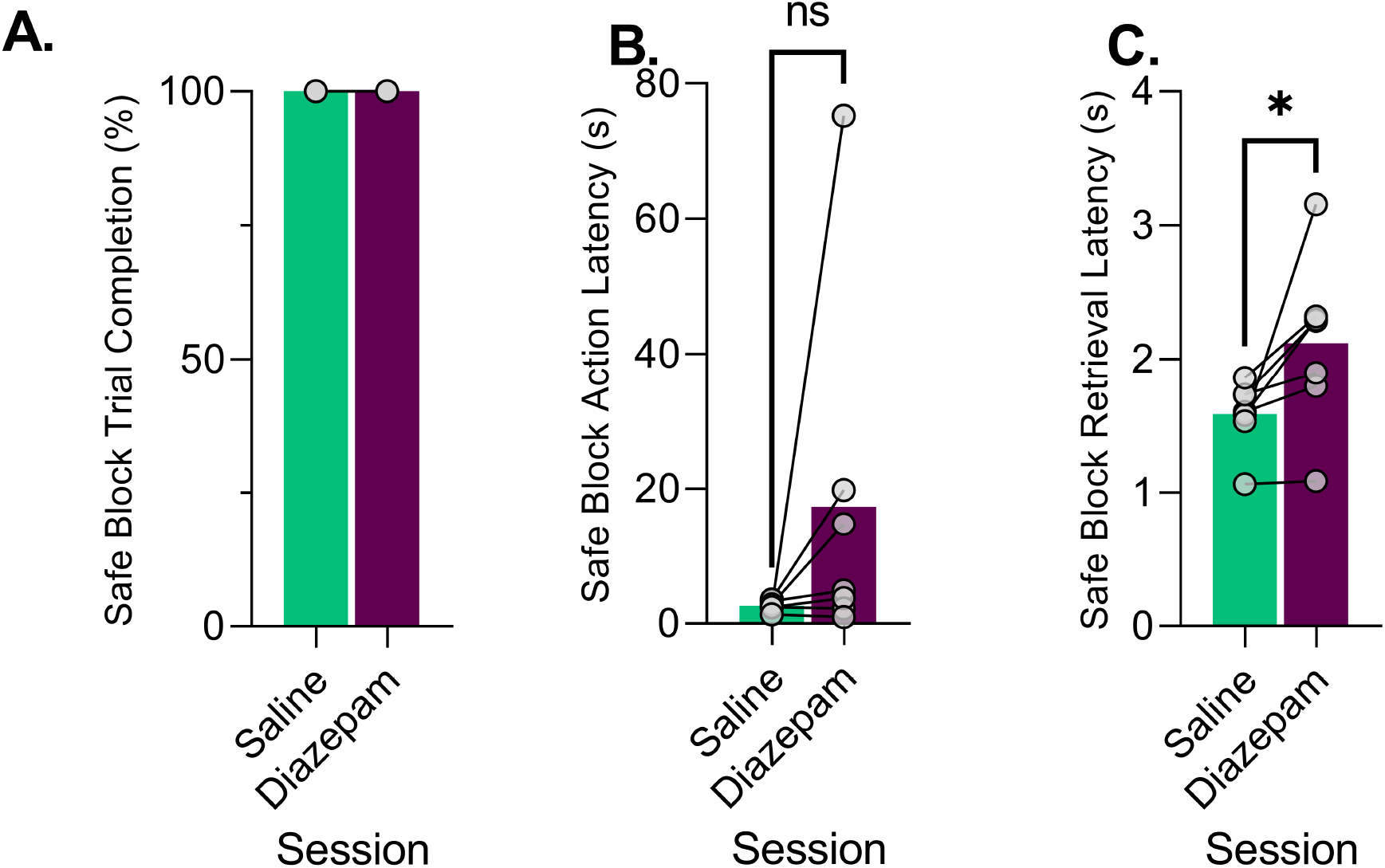
Average and individual (grey dots) behavior during the safe (0% risk) block for saline and diazepam pretreatment sessions. **A**. Percentage of completed trials in the safe block. **B**. Latency to perform the action for the safe block. C. Latency to retrieve the food pellet in the safe block. Green - Saline, Magenta - Diazepam. * p*<*.05, ns= not significant. Back to results.

**Supp. Figure 9:**
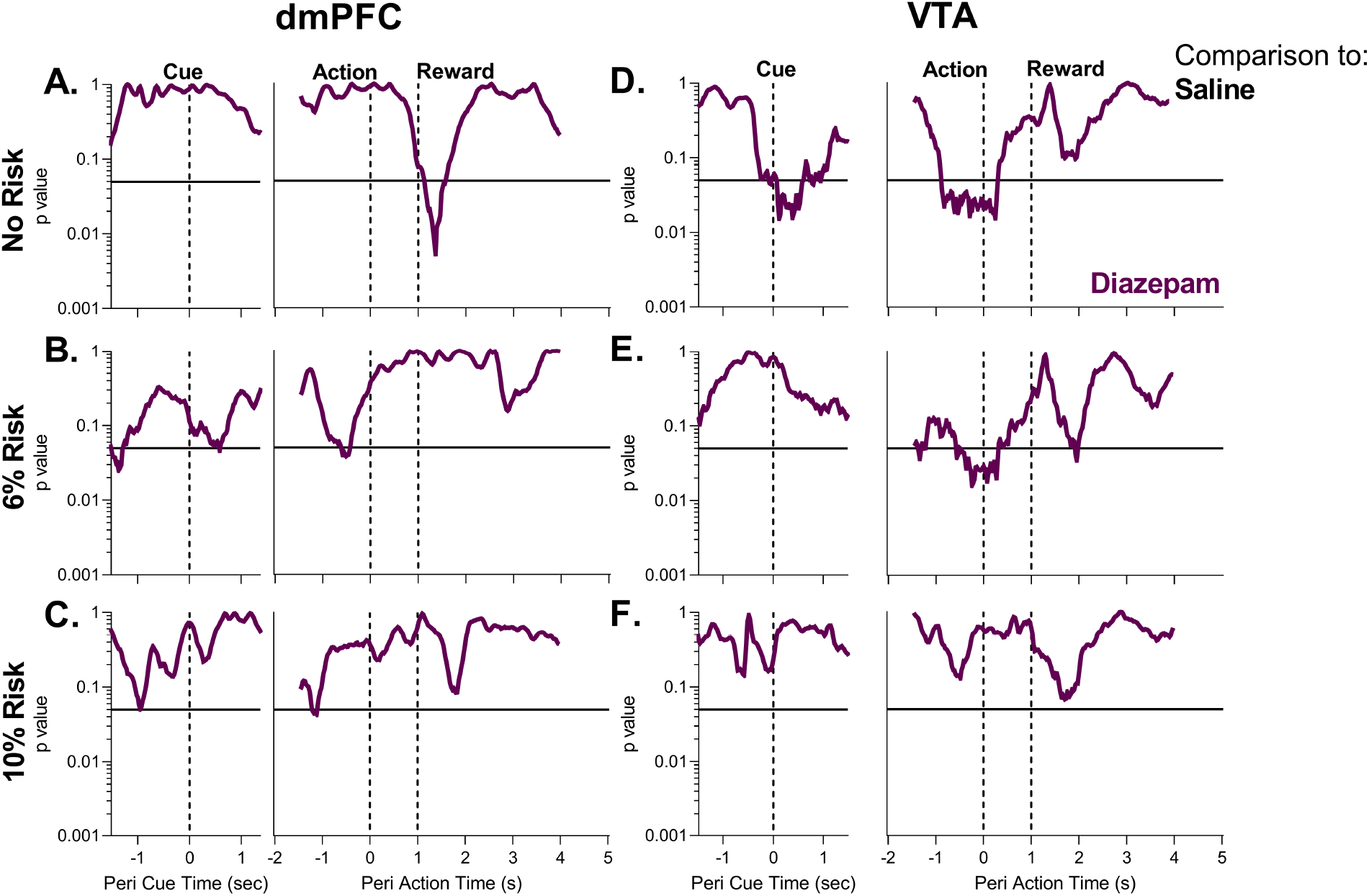
Permutation test results for recordings in the dmPFC and VTA for unpunished PRT trials after saline or diazepam (2 mg/kg) pretreatment. **A-C**. dmPFC p-value for tests in the cue, action, and reward delivery periods for each block during the PRT task. **D-F**. VTA p-value for tests in the cue, action, and reward delivery periods for each block during the PRT task. All comparisons are to the Saline session. Magenta – Diazepam. Solid black line indicates p =.05.Back to results.

**Supp. Figure 10:**
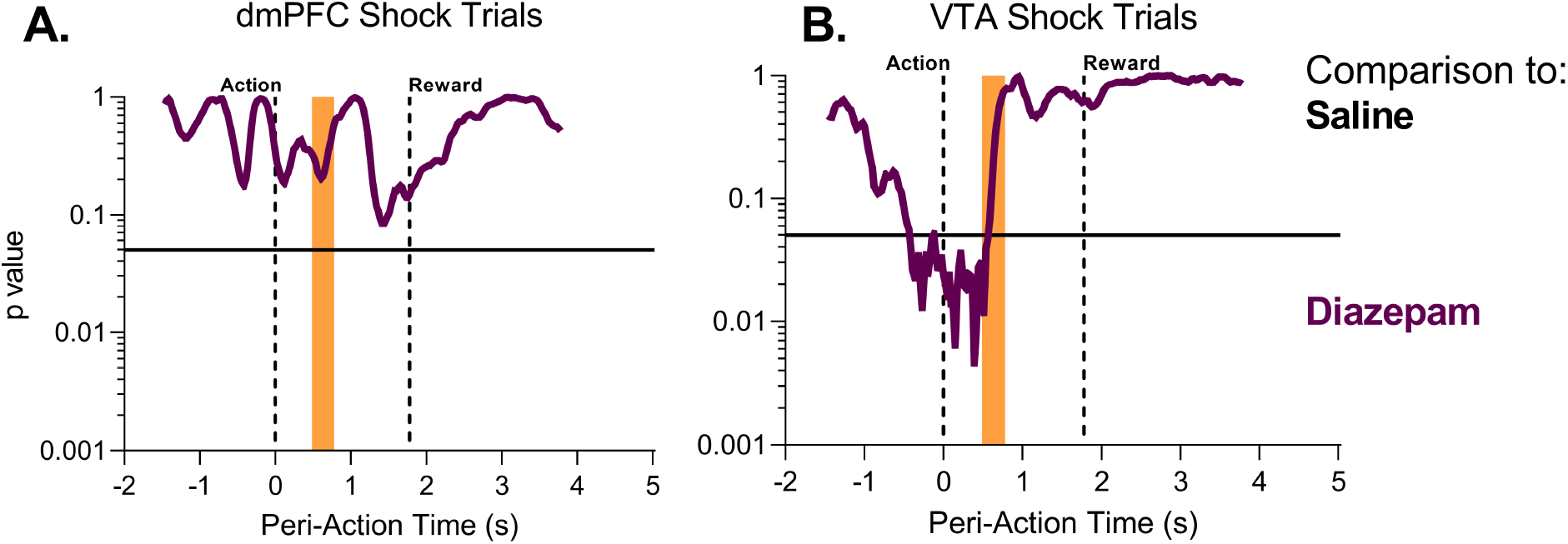
Permutation test results for recordings in the dmPFC and VTA for punished PRT trials after diazepam (2 mg/kg) pretreatment. **A**. dmPFC p-value for tests in the action, footshock (orange bar), and reward delivery periods for the first three session in the PRT task. **B**. VTA p-value for tests in the action, footshock (orange bar), and reward delivery periods for the first three session in the PRT task. All comparisons are to the Saline session. Magenta – Diazepam. Solid black line indicates p =.05.Back to results.

**Supp. Figure 11:**
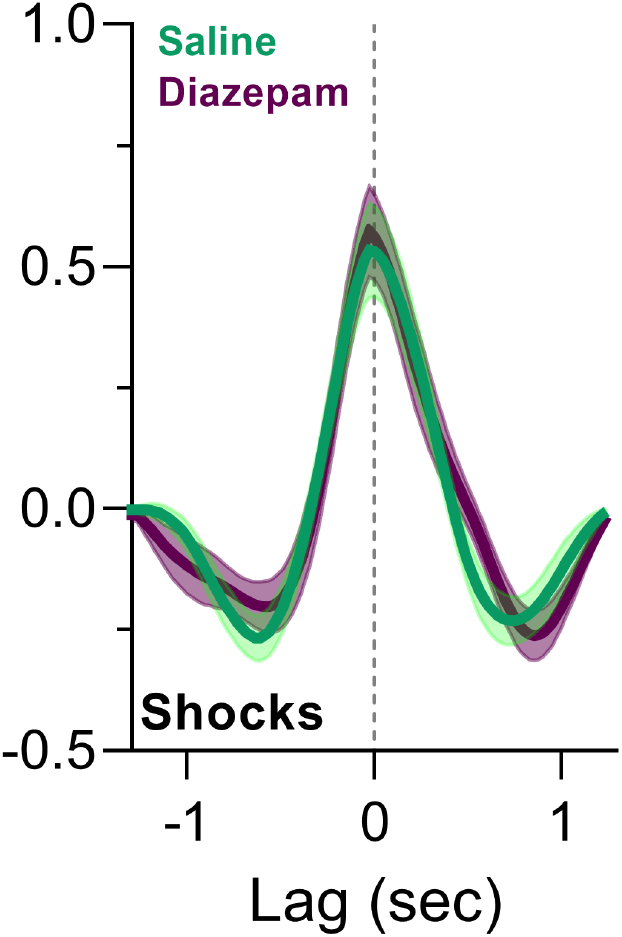
Cross-correlation results for dmPFC-VTA correlated activity for punished (footshock) trials after Saline (green) or Diazepam (magenta) pretreatment. Back to results.

